# Comparative genomics of *Cadophora luteo-olivacea* reveals a divergent lineage, conserved functional repertoires, and strain-level variation in pathogenicity

**DOI:** 10.64898/2026.04.07.716880

**Authors:** C. Leal, R. Bujanda, A. Eichmeier, J. Pečenka, E. Hakalova, L. Antonielli, S. Compant, D. Gramaje

## Abstract

*Cadophora luteo-olivacea* is an ecologically versatile fungus associated with grapevine trunk diseases, yet the extent to which strains from different hosts and environments differ in genome composition, functional potential, and pathogenicity remains poorly understood. Here, we performed a comparative genomic analysis of 12 *C. luteo-olivacea* isolates recovered from grapevine, almond, apple, *Crocus* bulbs, soil, air, wastewater, and deep-sea sediment. Genome assemblies were highly complete (BUSCO >99%) and ranged from 46.94 to 50.70 Mbp. Pairwise average nucleotide identity (ANI) revealed a cohesive 11-strain group and one markedly divergent strain, CBS 266.93. Phylogenomic analysis based on 2,645 single-copy orthologs further showed that CBS 266.93 lies outside the main *C. luteo-olivacea* clade and forms a sister relationship with *Cadophora malorum*, indicating that its taxonomic placement warrants reassessment. Across the remaining strains, broad functional conservation was observed, including similar KOG profiles, extensive carbohydrate-active enzyme repertoires (798-849 genes per genome), and abundant biosynthetic gene clusters (26-35 per genome). Transposable element content varied substantially among strains (0.67-4.45% of genome), but this variation did not parallel overall functional profiles. All isolates colonized grapevine leaves *in vitro*, although lesion severity differed significantly among strains, indicating conserved plant-colonizing capacity with quantitative variation in aggressiveness. Small RNA profiling of inoculated grapevine leaves further revealed isolate-associated differences in host miRNA family expression, particularly for miR398, miR827, and miR156. Together, these results show that most *C. luteo-olivacea* strains share a conserved genomic framework compatible with plant colonization, while retaining lineage-and strain-level phenotypic and host-associated variation.

## INTRODUCTION

The genus *Cadophora* Lagerberg & Melin was established in 1927 with *Cadophora fastigiata* as the type species (Lagerberg et al., 1927), and currently includes more than 30 described taxa with broad ecological and biological diversity (Chen et al., 2022). Species of *Cadophora* have been recovered from a wide range of substrates and environments, including decaying wood, soils, water-associated habitats, and plant tissues, highlighting the ecological breadth of the genus. Among them, *Cadophora luteo-olivacea* has attracted particular attention because of its recurrent association with economically important crops, including grapevine (Gramaje et al., 2018; Travadon et al., 2022), soybean (Allington & Chamberlain, 1948), and kiwifruit (Prodi et al., 2008).

*Cadophora luteo-olivacea* is notable for its apparent ecological versatility. It has been reported from decaying wood (Nilsson, 1973; Morrell & Zabel, 1985; Blanchette et al., 2004; Held et al., 2005; Arenz et al., 2006), soils (Kerry, 1990; Aislabie et al., 2001; Arenz et al., 2006; Hujslová et al., 2010), and marine or deep-sea environments (Marchese et al., 2021), but also from diseased plant tissues. This combination of environmental persistence and plant association suggests a flexible lifestyle in which saprotrophic and pathogenic capacities may coexist within the same species. Understanding how this versatility is reflected at the genome level remains an important question for fungal biology, especially in taxa that occupy the interface between environmental and plant-associated niches. More broadly, comparative analyses across plant ascomycete pathogens indicate that shifts in lifestyle are often accompanied by distinctive balances of CAZyme repertoires, secondary metabolism, and other virulence-associated functions, underscoring the value of placing *C. luteo-olivacea* within a wider comparative framework (Wang et al., 2022).

In viticulture, *C. luteo-olivacea* is best known for its association with grapevine trunk diseases, particularly Petri disease and Esca (Gramaje et al., 2018). These diseases are linked to vascular dysfunction, wood discoloration, necrosis, and long-term decline in vine productivity (Gramaje & Eichmeier, 2026). Previous pathogenicity studies have shown that *C. luteo-olivacea* can induce wood lesions and vascular symptoms in grapevine, supporting its role as a genuine pathogen rather than merely a secondary colonizer (Halleen et al., 2007; Gramaje et al., 2011; Úrbez-Torres et al., 2014; Travadon et al., 2015). Nevertheless, the genomic basis of its pathogenic potential, and the extent to which strains from diverse ecological origins differ in their colonization-related repertoires, remain poorly resolved.

Earlier population-level studies provided initial evidence of diversity within *C. luteo-olivacea*. In particular, Gramaje et al. (2011, 2014) identified genetic structure within grapevine-associated populations and reported distinct clonal lineages in Spanish and South African isolates. However, these studies were based on the marker systems available at the time and could not resolve genome-wide diversity or broader patterns of functional conservation. Comparative genomics now offers the opportunity to revisit these questions at much higher resolution, allowing the simultaneous assessment of lineage divergence, repeat content, biosynthetic potential, and candidate functions linked to plant colonization and environmental persistence (Rouxel & Balesdent, 2017; Sánchez-Vallet et al., 2018). Comparable genome-wide studies in other grapevine trunk pathogens have further shown that gene families associated with plant cell wall degradation, secondary metabolism, and nutrient uptake can evolve differentially among lineages, highlighting the value of extending this type of analysis to *Cadophora* (Morales-Cruz et al., 2015).

Here, we conducted a comparative genomic analysis of 12 *C. luteo-olivacea* strains isolated from different hosts and environments, including grapevine, almond, apple, *Crocus* bulbs, soil, air, wastewater, and deep-sea sediment. Our objectives were to: (i) characterize genome-wide diversity among strains, (ii) determine whether broad functional repertoires are conserved across ecologically diverse isolates, (iii) identify lineage-level variation in features such as transposable elements, carbohydrate-active enzymes, and biosynthetic gene clusters, and (iv) compare these genomic patterns with quantitative pathogenicity assays and host-associated small RNA profiles in grapevine. By combining comparative genomics with phenotypic and host-response data, this study provides a broader framework for understanding diversity, conservation, and plant-associated behaviour in *C. luteo-olivacea*.

## MATERIALS AND METHODS

### Fungal Strains and Culture Collection

Twelve *C. luteo-olivacea* isolates representing diverse ecological origins were selected for genomic analysis (Table 1). The collection included strains isolated from diseased grapevines and almonds, as well as reference isolates obtained from the Centraalbureau voor Schimmelcultures (CBS), Netherlands. CBS isolates originated from various plant hosts including *Malus sylvestris* and *Crocus* bulbs, and diverse environments including deep-sea sediment, wheat field soil, air, and wastewater, providing comprehensive representation of the species’ ecological range.

### DNA Isolation and Fungal Identification

Fungal strains were cultured on potato dextrose agar (PDA; Conda Laboratories, Madrid, Spain) for 10 days at room temperature under standard laboratory conditions. Mycelium was harvested using sterile scalpels and ground to fine powder in liquid nitrogen using mortar and pestle. DNA extraction was performed from 100 mg of powdered mycelium using the DNeasy Plant Mini Kit (Qiagen, Hilden, Germany) following manufacturer protocols. Extracted DNA was purified and concentrated using Amicon Ultra-0.5 mL Centrifugal Filters (30 kDa cutoff; Millipore-Merck, Bedford, MA, USA) to remove potential PCR inhibitors and ensure high-quality template for downstream applications.

Species identification was confirmed by sequencing the beta-tubulin TUB2 gene, identified as the most informative locus for *Cadophora* species delimitation (Gramaje et al., 2025), following established protocols (Travadon et al., 2022). PCR amplification and sequence analysis were performed according to standardized procedures, with resulting sequences subjected to phylogenetic analysis using Maximum Likelihood methods implemented in MEGA v.6 (Tamura et al., 2013). The optimal evolutionary model was selected using the Bayesian Information Criterion in jModelTest 2.1.10 (Darriba et al., 2012), and branch support was assessed using 1,000 bootstrap replicates. *Hyaloscypha finlandica* isolates served as outgroups for phylogenetic reconstruction.

### Genome Sequencing and Assembly

High-throughput sequencing was performed using Illumina NextSeq 500 platform with V2 reagent kit (2 × 150 bp paired-end reads), generating approximately 50 million read pairs per sample (LGC Genomics GmbH, Berlin, Germany). Raw sequence data underwent comprehensive quality control including PhiX contamination assessment using Bowtie 2 v2.3.4.3 (Langmead & Salzberg, 2012) and adapter removal using fastp v0.19.5 (Chen et al., 2018). Sequence quality metrics and length distributions were evaluated using FastQC (Andrews, 2010) to ensure data suitability for genome assembly.

*De novo* genome assembly was conducted using SPAdes v3.14.0 (Bankevich et al., 2012) with optimized parameters for fungal genomes. Low-coverage contigs (<2× depth) were filtered to remove potential assembly artifacts. Contamination screening was performed using BlobTools (Laetsch & Blaxter, 2017) to identify and remove non-fungal sequences. Assembly quality was assessed using QualiMap v2.2 (Okonechnikov et al., 2016) and QUAST v5.0.0 (Gurevich et al., 2013), while genome completeness was evaluated using BUSCO v4.0.6 (Simão et al., 2015) against the Ascomycota gene set.

Taxonomic verification was performed by extracting complete Internal Transcribed Spacer (ITS) regions using ITSx v1.1 (Bengtsson-Palme et al., 2013) and comparing sequences against the UNITE database (Nilsson et al., 2019) using BLAST searches to confirm species identity and detect potential misidentifications.

### Gene Prediction and Genome Annotation

Comprehensive gene prediction was performed using the Funannotate pipeline (Palmer & Stajich, 2020) with multiple evidence sources. Genome contigs were preprocessed using minimap2 (Li, 2018) to identify and remove redundant sequences, followed by soft-masking of repetitive elements using tantan (Frith, 2011). Gene prediction employed a consensus approach combining multiple algorithms including GeneMark-ES (Borodovsky & Lomsadze, 2011), SNAP (Korf, 2004), GlimmerHMM (Majoros et al., 2004), and modified BUSCO (Simão et al., 2015), with results integrated using EvidenceModeler (Haas et al., 2008) to generate consensus gene models.

Protein evidence from related fungi was aligned to genomic contigs using Diamond (Buchfink et al., 2015) and Exonerate (Slater & Birney, 2005) to provide external validation for gene predictions. High-quality Augustus predictions were generated following iterative training on the consensus gene set (Stanke et al., 2006). Transfer RNA genes were identified using tRNAscan-SE (Lowe & Eddy, 1997) to complete the structural annotation.

### Functional Annotation of Predicted Genes

Comprehensive functional annotation was performed using multiple complementary approaches: (1) eggNOG-mapper v2 (Huerta-Cepas et al., 2019) against the latest eggNOG database for ortholog assignment and functional classification, (2) InterProScan 5 (Jones et al., 2014) for protein domain identification, (3) Diamond BLASTP searches against UniProt database (The UniProt Consortium, 2019) for homology-based annotation, and (4) KOG (Clusters of Orthologous Groups) analysis for comparative functional genomics. Results were integrated to provide comprehensive functional profiles for each genome.

### Transposable Element Analysis

Transposable element identification and classification were performed using the EDTA (Extensive de novo TE Annotator) pipeline (Ou et al., 2019) implemented in a local Conda environment. This comprehensive approach identified and classified major TE families including LTR retrotransposons (Copia, Gypsy), DNA transposons (CACTA, Mutator, Pif-Harbinger, Tc1-mariner, Hat), and non-LTR elements. TE content and distribution patterns were analysed to assess their potential impact on genome evolution and gene regulation.

### Carbohydrate-Active Enzyme Analysis

Carbohydrate-active enzymes (CAZymes) were identified and classified using the dbCAN2 HMM-based system (Zhang et al., 2018) with stringent parameters (E-value < 1×10 ¹, coverage > 0.35). All CAZyme classes were analyzed including Glycoside Hydrolases (GH), Carbohydrate Esterases (CE), Glycosyl Transferases (GT), Polysaccharide Lyases (PL), Auxiliary Activities (AA), and Carbohydrate-Binding Modules (CBM). Secreted CAZymes were predicted using signal peptide prediction algorithms to identify enzymes likely involved in plant cell wall degradation.

Comparative CAZyme analysis included reference genomes from fungi with diverse lifestyles including root pathogens, aerial pathogens, trunk pathogens, white-rot degraders, indoor biodeterioration agents, saprotrophs, dark septate endophytes, and mycorrhizal symbionts to contextualize *C. luteo-olivacea* CAZyme profiles within broader fungal ecology.

### Secondary Metabolite Cluster Analysis

Biosynthetic gene clusters for secondary metabolites were predicted using antiSMASH fungal version (Blin et al., 2019) with default parameters. Identified clusters were classified into major categories including polyketide synthases (PKS), non-ribosomal peptide synthetases (NRPS), terpene synthases, and hybrid clusters. Cluster completeness and similarity to known metabolites were assessed to predict potential bioactive compounds and their roles in fungal ecology and pathogenicity.

### Comparative Genomic Analysis

Genome-wide comparisons were performed among the 12 *C. luteo-olivacea* genomes generated in this study. Pairwise average nucleotide identity (ANI) values were calculated using fastANI v1.33 (Jain et al., 2018) with default parameters. ANI values were computed for all pairwise combinations of the 12 *C. luteo-olivacea* genomes and visualized as a heatmap. The resulting ANI matrix was used to assess genome-wide relatedness among strains and to identify divergent lineages within the dataset. Because ANI thresholds commonly used for species delimitation were developed primarily for prokaryotes, ANI values in this study were interpreted comparatively and together with phylogenomic evidence, rather than as a standalone criterion for fungal species delimitation. Metadata on host or source, environment, geographic origin, and melanized phenotype were integrated into the heatmap for interpretative purposes.

To further assess the phylogenomic placement of the analysed strains, a supplementary phylogenomic analysis was performed using single-copy orthologs. In addition to the 12 *C. luteo-olivacea* genomes, three reference genomes were included: *Cadophora malorum*, *Oidiodendron maius* Zn, and *Botrytis cinerea* B05.10. *O. maius* and *B. cinerea* were included as external reference taxa, whereas *C. malorum* was used to evaluate the placement of divergent lineages within the genus. Single-copy orthologs were identified using BUSCO v6.0.0 in genome mode with the *ascomycota_odb12* lineage dataset and Miniprot-based gene prediction. Orthologs present as single-copy genes across all selected genomes were extracted and aligned individually with MAFFT. Poorly aligned regions were removed prior to concatenation. The resulting filtered alignments were concatenated into a supermatrix, and maximum-likelihood phylogenetic reconstruction was performed with IQ-TREE using automatic model selection. Node support was assessed with 1,000 SH-aLRT replicates and 1,000 ultrafast bootstrap replicates. The resulting tree was used as supplementary evidence to refine the interpretation of ANI-based relationships among strains (Figure S1). Because CBS 128.578 and CBS 141.41 showed ANI = 100% and their provenance metadata could not be independently clarified, comparisons involving these two accessions were interpreted cautiously as potentially redundant genomic sampling. Both strains were retained in the ANI and phylogenomic analyses for transparency.

### Pathogenicity Testing

#### Plant Material and Cultivation

Pathogenicity assays were conducted using *Vitis vinifera* cv. Chardonnay clone CHAR PO-156/4 maintained under controlled *in vitro* conditions. Shoot samples collected at the end of the 2024 growing season were processed for micropropagation using established protocols (Baránek et al., 2010). Nodal segments were cultured on Murashige and Skoog (MS) medium supplemented with 1.33 μM 6-benzylaminopurine (BA) and 0.57 μM indole-3-acetic acid (IAA) under controlled environmental conditions (23°C, 16/8 h photoperiod). Following three weeks of growth, plantlets were transferred to fresh medium in individual vessels. Rooting was induced on MS medium containing 0.81 μM naphthaleneacetic acid (NAA) after six weeks of vegetative growth.

#### Plant Inoculation Protocol

Pathogenicity tests were performed on six-week-old rooted plantlets maintained under controlled growth chamber conditions (23°C, 16 h light/8 h dark photoperiod). Single leaves per plant were inoculated with 3-mm agar plugs taken from the actively growing colony margin of 10-day-old *C. luteo-olivacea* cultures. Plugs were placed directly onto the non-wounded leaf surface and left in contact without additional fixation. Control plants received sterile PDA plugs using the same procedure. Disease severity was quantified from digital images using Fiji/ImageJ at 3, 5, 7, and 10 days after inoculation (DAI). Each treatment included five biological replicates (plants), each with two inoculated leaves, and the experiment was repeated independently after two weeks.

#### Statistical analysis of pathogenicity assays

Lesion severity at 10 days after inoculation (DAI) was analysed using a linear model with strain and replicate as fixed effects. For each plant, lesion values from the two inoculated leaves were averaged prior to analysis. Pairwise comparisons among strains were performed using Tukey’s post hoc test.

### RNA Extraction and miRNA Analysis

#### Sample Collection and RNA Isolation

Leaf tissue was collected at 10 DAI from the whole inoculated leaves, immediately frozen in liquid nitrogen, and stored at −80°C until RNA extraction. Total RNA was extracted using PureLink™ Plant RNA Reagent (Thermo Fisher Scientific) according to the manufacturer’s instructions. RNA quality and integrity were assessed using an Agilent 2100 Bioanalyzer with the RNA 6000 Nano Kit (Agilent Technologies), and samples with RNA Integrity Number (RIN) values below 7 were excluded. RNA concentration was determined using a Modulus™ Single Tube Multimode Reader (Turner Biosystems) with the Quant-iT™ RNA Assay Kit. Only samples with concentrations ≥ 5 ng μL ¹ were retained and normalized to 5 ng μL¹.

#### Library Preparation and Sequencing

Small RNA libraries were prepared using the QIAseq miRNA Library Kit (Qiagen) following the manufacturer’s protocol. Library quality and concentration were assessed using the Agilent High Sensitivity DNA Kit and additional fluorometric and qPCR-based quantification methods (MCNext™ SYBR® Fast qPCR Library Quantification Kit, MCLAB) on a Rotor-Gene 3000 system. Libraries were pooled to a final concentration of 2 nM, assuming an average insert size of 180 bp. Sequencing was performed on an Illumina MiniSeq platform using the High Output Reagent Kit (75-cycle), generating 36-nt single-end reads. Three biological replicates were sequenced for each fungal isolate.

#### Small RNA Processing, miRNA Annotation, and Differential Expression Analysis

Raw small RNA reads were quality filtered (Phred score ≥ Q30), adapter trimmed, and collapsed into unique sequences. Sequences retained for downstream analysis were filtered by abundance, requiring at least 10 counts in at least three samples. Differential expression analysis was performed in DESeq2 using raw count data and a design formula of ∼ condition. Because control libraries showed substantially lower read depth than inoculated samples, formal statistical comparisons were restricted to differences among fungal isolates rather than isolate-versus-control contrasts. Within this framework, BV-144 was used as the reference level in the DESeq2 model to express log2 fold changes among inoculated treatments. This choice was analytical rather than biological and does not affect which miRNA families are identified as differentially represented, but only the direction and magnitude of fold-change estimates relative to that isolate.

To improve biological interpretability and account for isomiR variation, filtered sequences were annotated against miRBase Release 22.1 using a BLAST-like alignment approach without gaps and allowing up to two mismatches. Only high-confidence matches were retained. Sequences assigned to the same miRNA family were collapsed prior to family-level analysis. Variance-stabilizing transformation (VST) was used for exploratory data visualization, including principal component analysis. Differential representation of miRNA families was summarized as log2 fold change relative to BV-144, with significance assessed using Wald tests and Benjamini–Hochberg adjusted P values.

## RESULTS

### Genome Assembly and Quality Assessment

High-quality genome assemblies were obtained for all 12 *C. luteo-olivacea* strains, with genome sizes ranging from 46.94 to 50.70 Mbp (Table S1). All assemblies demonstrated exceptional completeness with BUSCO scores exceeding 99%, indicating successful capture of conserved single-copy orthologs typical of Ascomycete fungi. Contig numbers ranged from 155 to 603, reflecting varying degrees of assembly contiguity, while GC content remained consistently between 47-48% across all strains. Gene prediction identified 15,653 to 17,036 protein-coding genes per genome, consistent with typical Ascomycete gene numbers. Sequencing coverage exceeded 100× for most samples, ensuring robust genome assembly and accurate variant detection. The high quality and completeness of these assemblies provide a solid foundation for comprehensive comparative genomic analyses.

### Average nucleotide identity reveals a highly divergent strain within the *C. luteo-olivacea* dataset

Pairwise ANI values among the 12 *C. luteo-olivacea* genomes ranged from 89.43% to 100% (mean = 95.28% ± 3.42%; median = 96.33%) (Figure 1). Eleven strains formed a high-similarity group with pairwise ANI values generally above 95%, whereas strain CBS 266.93, isolated from deep-sea sediment, consistently showed markedly lower ANI values (∼89.4%) relative to all other genomes, indicating substantial genomic divergence within the dataset.

**Fig 1.**
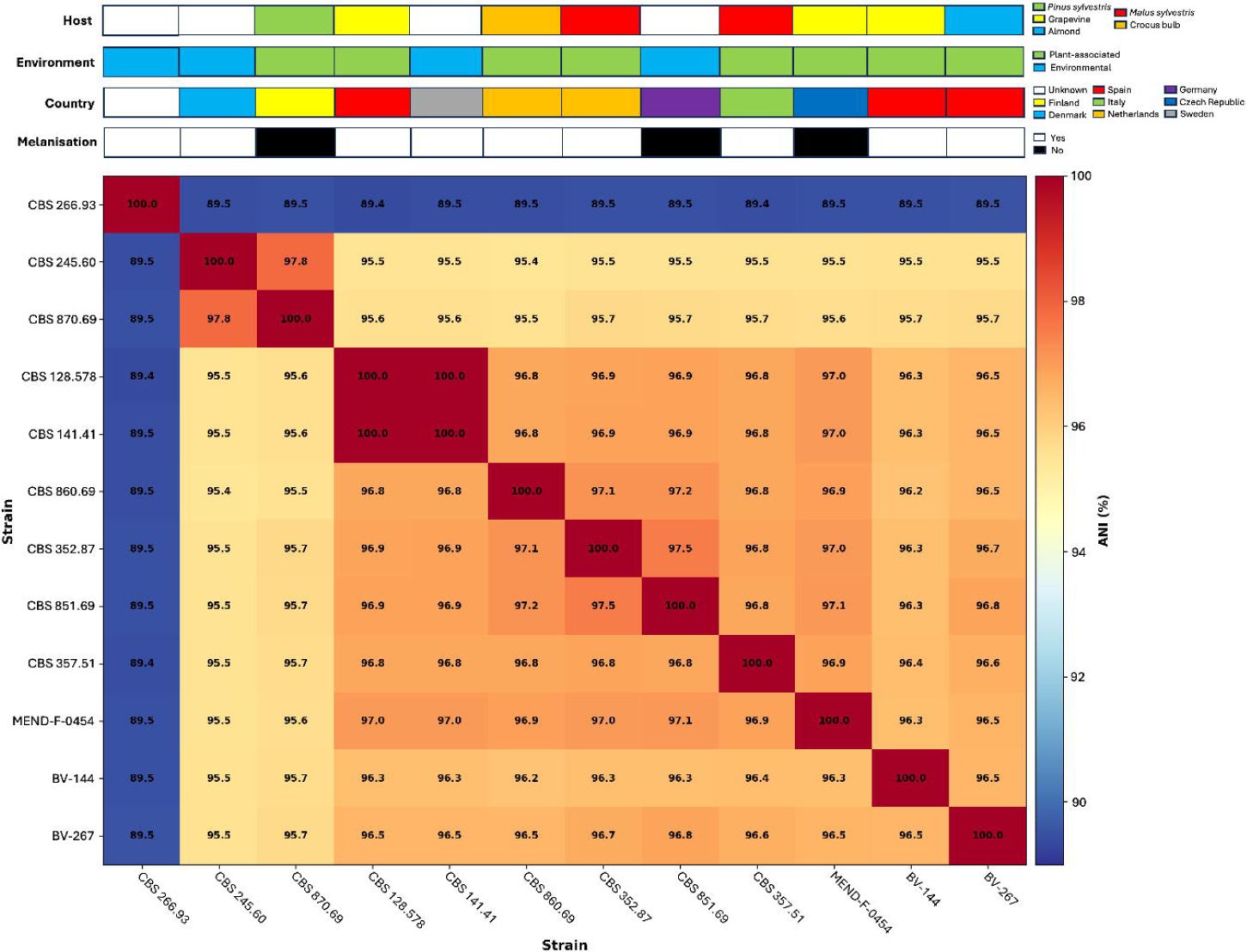

Within the main group, CBS 128.578 and CBS 141.41 showed ANI = 100%. Verification of the available metadata revealed inconsistencies in the reported provenance information that could not be resolved unambiguously from the accessible CBS records. We therefore retained both strains in the ANI heatmap and phylogenomic reconstruction for transparency, but interpreted downstream comparisons involving these two accessions with caution because they may represent redundant genomic sampling. Definitive resolution of whether this pattern reflects duplicate accessions, metadata inconsistency, or genuinely near-identical isolates would require raw-read-based SNP analysis.

Likewise, the melanized phenotype was not obviously associated with ANI-based genomic similarity, as the three melanized strains were distributed across different parts of the matrix. Overall, these results indicate a largely shared genomic background across most strains, together with the presence of one markedly divergent genome that warranted further phylogenomic investigation.

### Supplementary phylogenomic analysis supports the distinct placement of CBS 266.93

To further assess the position of CBS 266.93, a supplementary phylogenomic analysis was performed using 2,645 single-copy orthologs shared across the 12 *C. luteo-olivacea* genomes and three additional reference taxa (*Cadophora malorum*, *Oidiodendron maius*, and *Botrytis cinerea*) (Figure S1). Maximum-likelihood reconstruction based on the concatenated BUSCO supermatrix recovered a strongly supported clade containing 11 *C. luteo-olivacea* strains. In contrast, CBS 266.93 did not cluster within this main clade. Instead, it formed a sister relationship with *C. malorum*, with maximal nodal support (SH-aLRT/UFBoot = 100/100).

This topology indicates that CBS 266.93 is phylogenomically distinct from the main *C. luteo-olivacea* lineage represented by the other 11 strains. The supplementary phylogenomic result therefore refines the ANI-based interpretation by showing that the divergence of CBS 266.93 is not merely quantitative, but also reflected in its placement outside the principal *C. luteo-olivacea* clade.

### Functional Gene Content Analysis

KOG-based functional profiling revealed highly similar gene category distributions across all strains (Figure 2). Differences among strains were subtle and were observed mainly at the level of relative family abundance rather than broad functional composition. Overall, the heatmap showed limited functional differentiation among genomes, supporting the conservation of core cellular capacities despite the genomic divergence detected by ANI. Notably, CBS 266.93 did not show a strikingly distinct KOG profile relative to the remaining strains, suggesting that its phylogenomic divergence is not accompanied by major shifts in broad functional gene category abundance.

**Fig 2.**
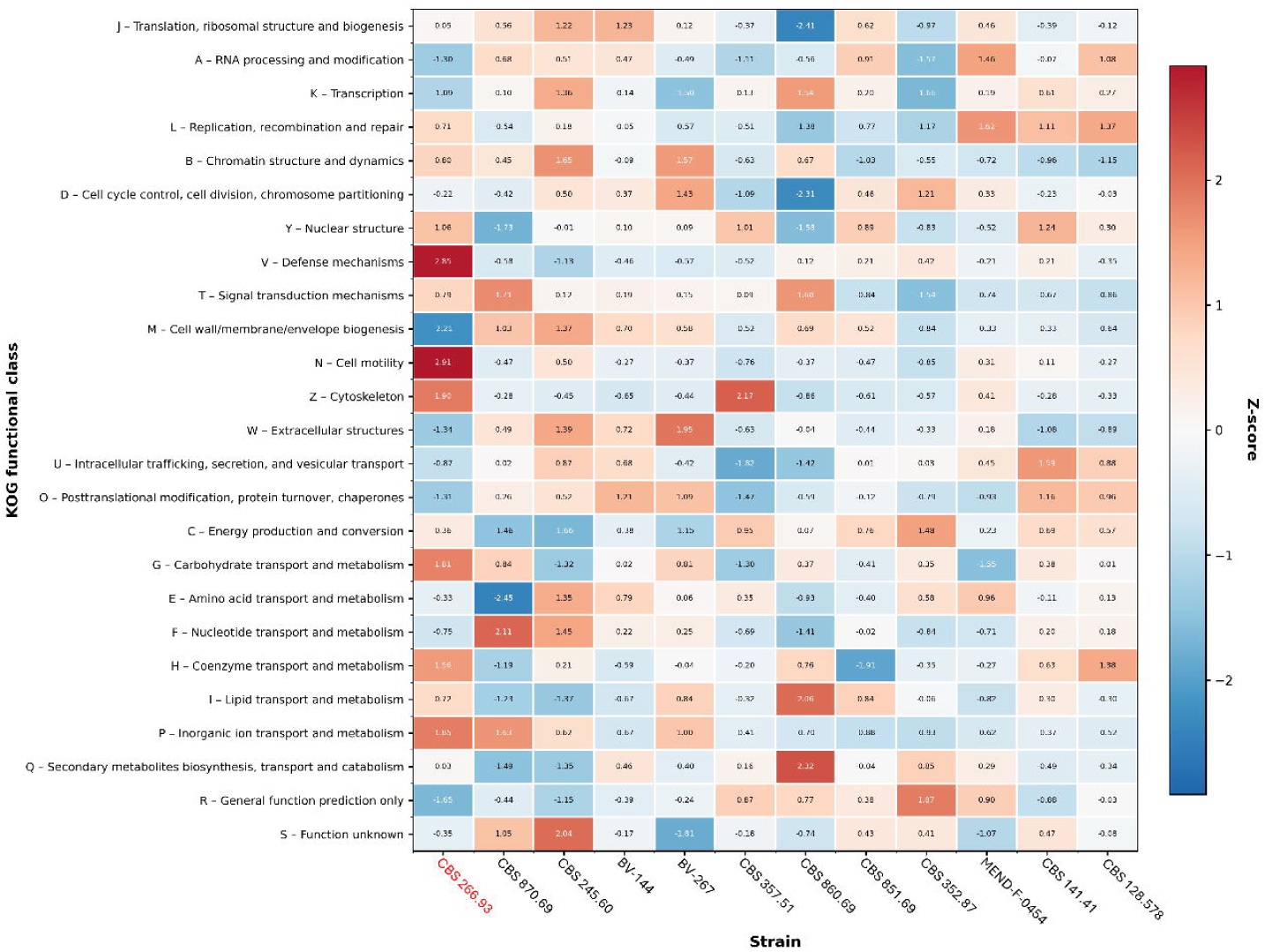

### Transposable Element Diversity

Transposable element content varied substantially among strains, with the proportion of genome masked by TEs ranging from 0.67% to 4.45% (Figure 3; Table S1). CBS 245.60 and CBS 870.69 showed the highest TE loads, whereas BV-144 and CBS 357.51 had the lowest values. CBS 266.93 showed an intermediate TE content (1.03%), indicating that its phylogenomic divergence was not accompanied by unusually high TE accumulation. Overall, these results suggest substantial variation in repeat content among strains despite their otherwise similar genome sizes and gene counts.

**Fig 3.**
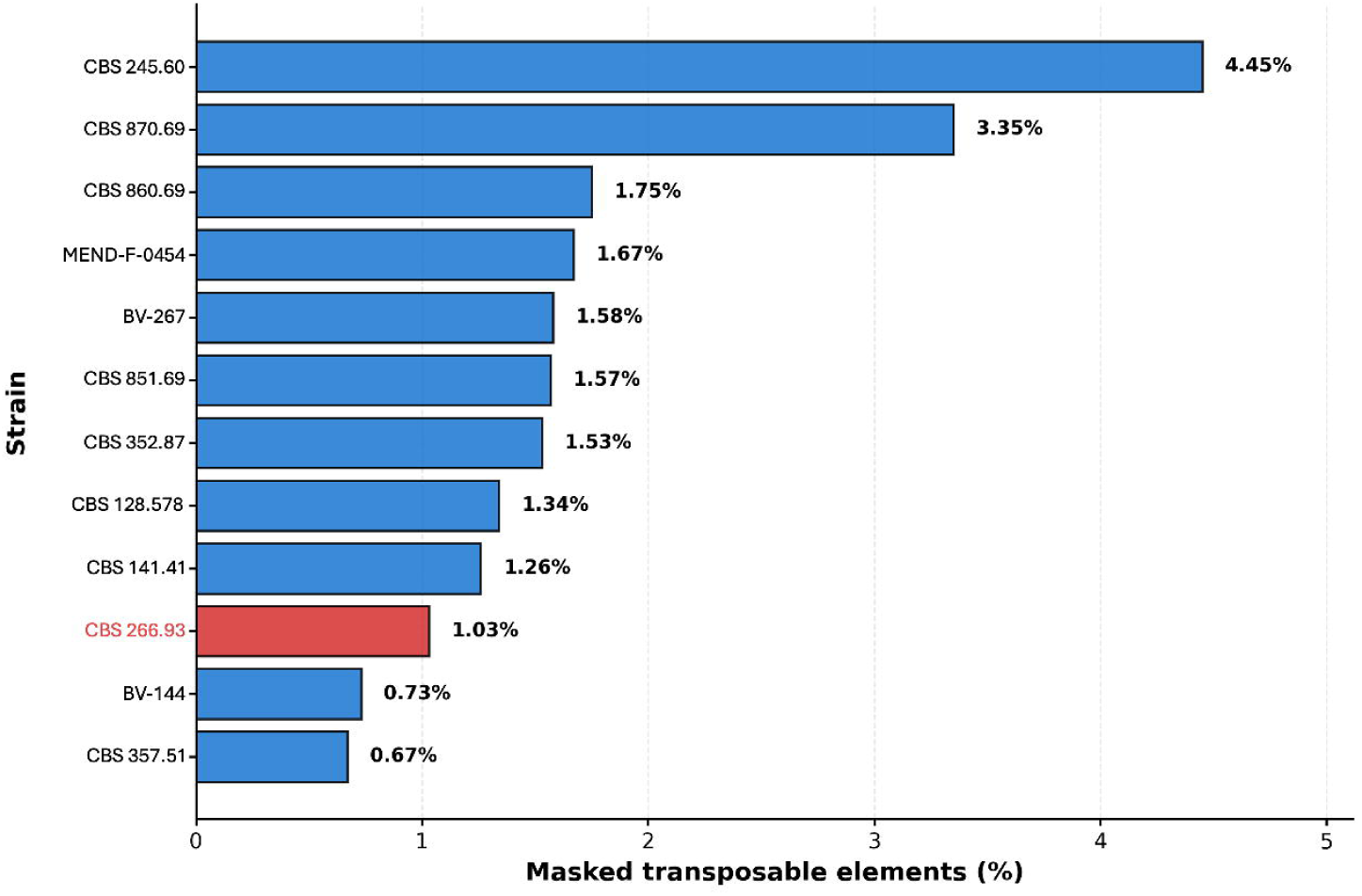

### Pathogenicity-Associated Genes

Analysis of selected pathogenicity-, metal resistance-, and ecology-related functions revealed broad conservation across the 12 *C. luteo-olivacea* strains (Figure 4; Table S2). Genes associated with necrosis-inducing proteins, proteases, lipases, and cytochrome P450s were detected in all strains, although copy number varied modestly among genomes. These results indicate that core functions potentially relevant to plant tissue colonization and pathogenicity are widely maintained across the species, irrespective of isolation source.

**Fig 4.**
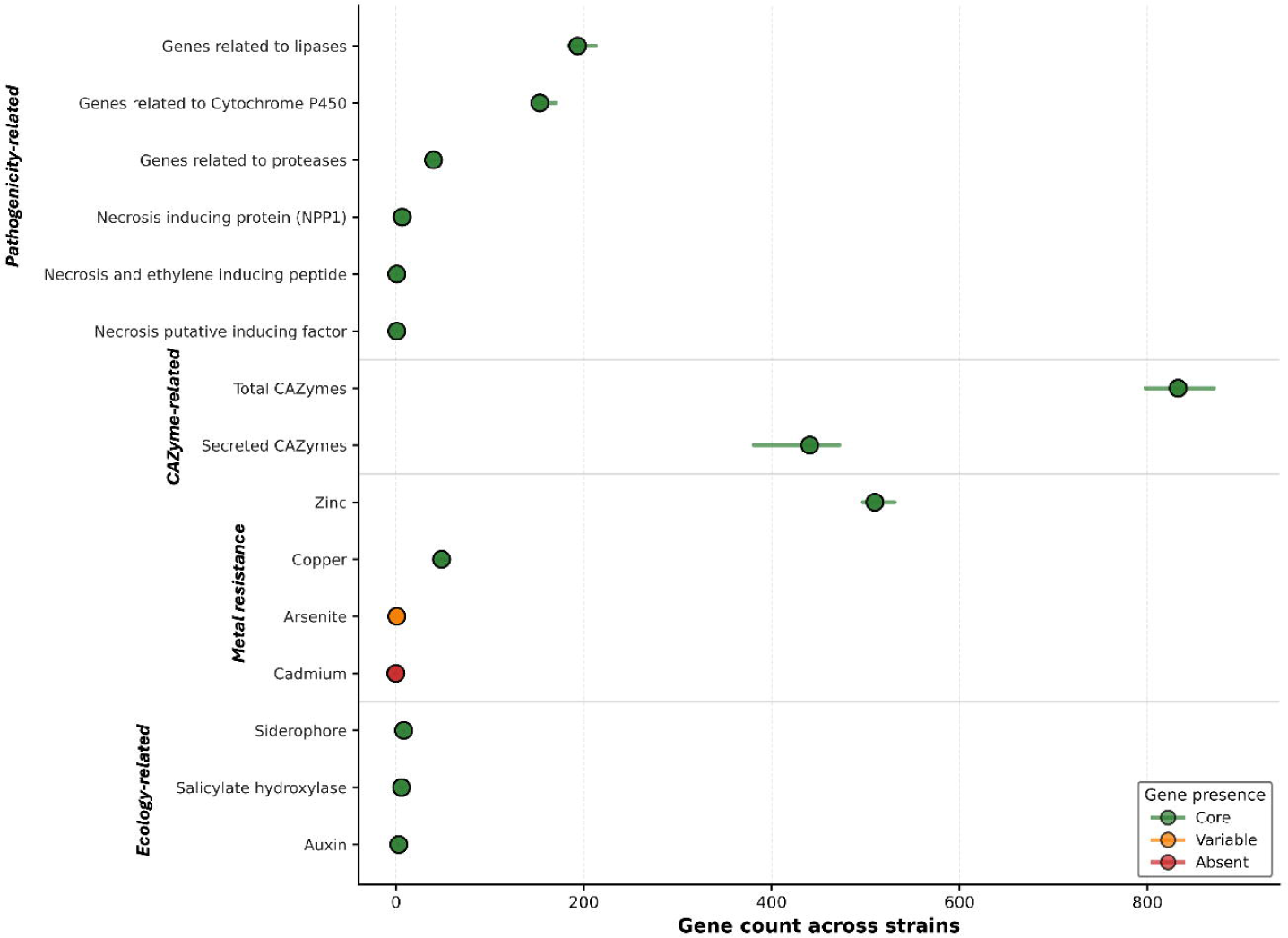

Metal and metalloid resistance-related genes were also broadly represented. Copper-and zinc-related genes were present in all strains, whereas arsenite-related genes varied among genomes. In contrast, cadmium resistance genes were not detected in any strain. This pattern suggests that tolerance to several environmentally relevant metals is conserved across the dataset, while other resistance-associated traits may be more unevenly distributed.

Additional functions of ecological and host-interaction relevance, including salicylate hydrolase, siderophore-related genes, and auxin-related genes, were also detected across the analysed genomes. Their widespread occurrence supports a shared capacity for nutrient acquisition, environmental persistence, and interaction with plant-associated niches. In particular, salicylate hydrolase is noteworthy because salicylic acid is a central component of plant defence signalling, and its degradation could potentially contribute to host colonization.

### Carbohydrate-Active Enzyme Arsenal

*Cadophora luteo-olivacea* strains possessed extensive CAZyme repertoires, ranging from 798 to 849 genes per genome (Figure 5; Table S3). Glycoside hydrolases were the most abundant class (388-420 genes), consistent with a strong capacity to degrade plant-derived polysaccharides. Between 380 and 450 CAZymes per strain were predicted to be secreted, indicating substantial extracellular enzymatic potential. Auxiliary activity enzymes were also abundant (162-173 genes per genome), including families associated with lignocellulose modification. Together, these features support a broad capacity for plant cell wall degradation and are consistent with the dual saprophytic and pathogenic lifestyle attributed to *C. luteo-olivacea*.

**Fig 5.**
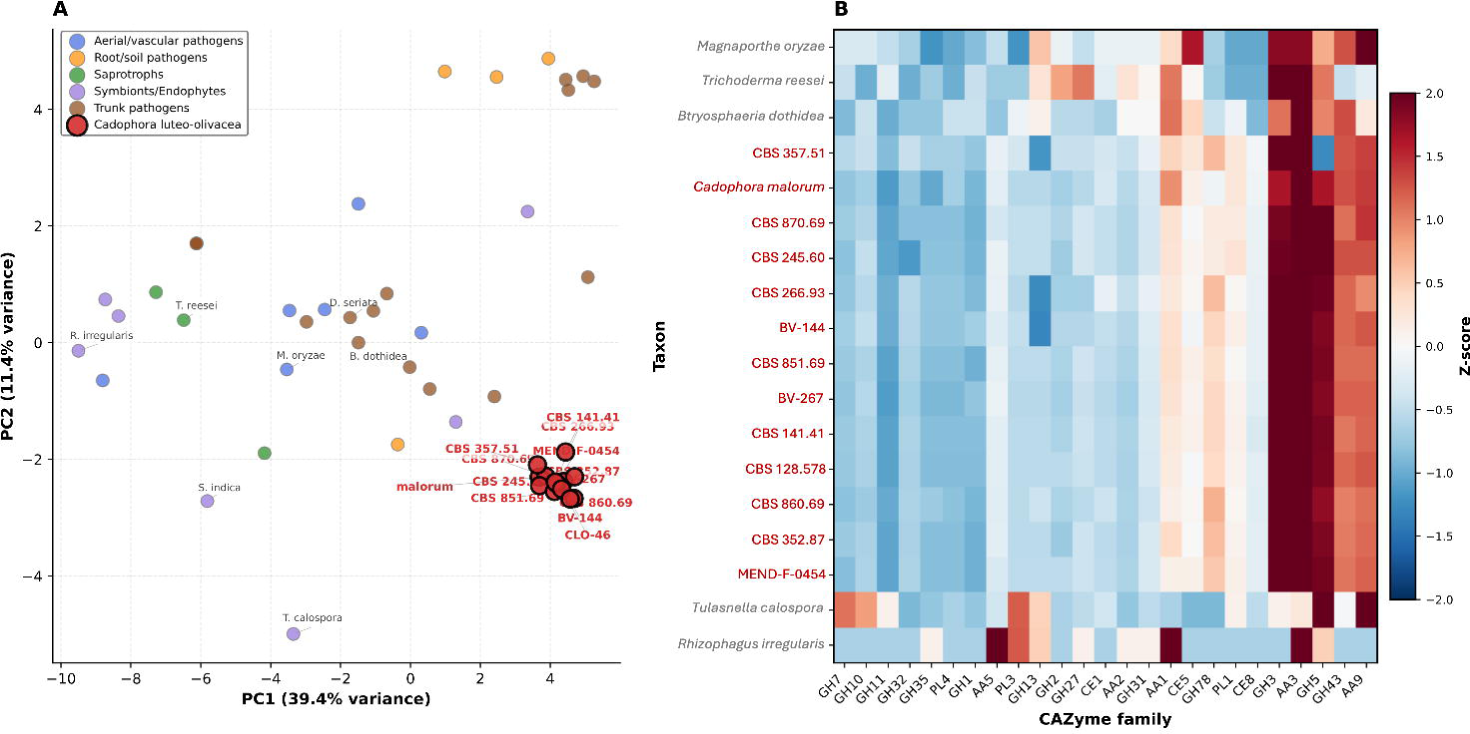

Comparative analysis further showed that *C. luteo-olivacea* CAZyme repertoires were broadly similar across strains and grouped close to selected grapevine trunk pathogens in multivariate space (Figure 5A). A heatmap of the most variable CAZyme families likewise indicated that, despite some variation in relative family abundance, the overall CAZyme composition remained highly conserved within *C. luteo-olivacea* (Figure 5B). This pattern suggests that the major enzymatic functions required for colonization of woody plant tissues are widely maintained across the species.

Among the families contributing most strongly to this profile were GH3, GH5, GH31, GH32, GH35, GH43, and GH78, all of which are associated with the degradation of major plant cell wall polysaccharides. Polysaccharide lyases and auxiliary activity families, including PL1, PL2, PL4, AA1, AA2, AA3, AA5, and AA9, were also well represented. Overall, these results indicate that *C. luteo-olivacea* shares a CAZyme repertoire structure consistent with other trunk-associated fungi while retaining a high degree of conservation across strains.

### Secondary Metabolite Biosynthetic Potential

Secondary metabolite biosynthetic gene clusters (BGCs) were abundant across all *Cadophora luteo-olivacea* genomes, with between 26 and 35 clusters predicted per strain (Figure 6). Terpene synthases, type I polyketide synthases (T1PKS), NRPS-like clusters, and non-ribosomal peptide synthetases (NRPS) represented the dominant BGC classes across the dataset. Hybrid clusters and minor classes such as indole-related clusters and other cluster types were present at lower frequencies.

**Fig 6.**
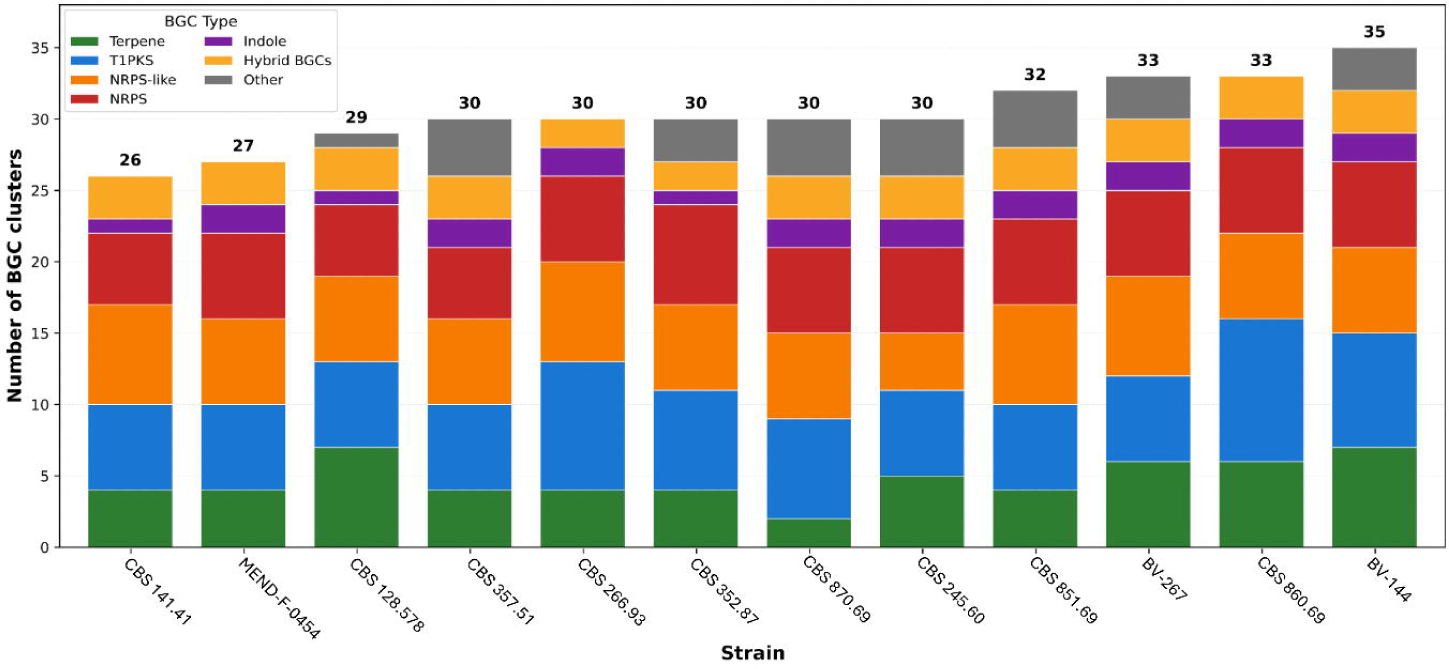

Although the total number of BGCs varied moderately among strains, their overall composition was broadly similar across the species (Figure 6). This pattern suggests that the major secondary metabolite biosynthetic capacities are largely conserved within *C. luteo-olivacea*. The presence of multiple terpene, PKS, and NRPS clusters indicates the potential for the production of diverse bioactive metabolites, including compounds potentially involved in fungal competition, environmental adaptation, or host interaction.

Most predicted clusters showed little or no similarity to characterized biosynthetic pathways in current databases, suggesting that *C. luteo-olivacea* may produce a substantial repertoire of previously undescribed secondary metabolites. Together, these results highlight the extensive biosynthetic potential of this species and suggest that secondary metabolites may contribute to its ecological versatility and ability to colonize diverse plant-associated environments.

### Pathogenicity Testing Results

*In vitro* pathogenicity assays showed that all *C. luteo-olivacea* strains were able to cause lesions on grapevine leaves, but lesion severity at 10 DAI differed significantly among strains (Figure 7; Table S4). Analysis of necrotic leaf area revealed a strong effect of strain, whereas no significant effect of experimental replicate was detected. BV-144 and CBS 128.578 produced the most extensive lesions, whereas CBS 851.69, CBS 352.87, and CBS 357.51 consistently showed the lowest lesion severity. Visual inspection also revealed contrasting mycelial phenotypes, with CBS 851.69, CBS 870.69, and MEND-F-0454 producing dark melanized growth, whereas most other strains produced greyish-white mycelium.

**Fig 7.**
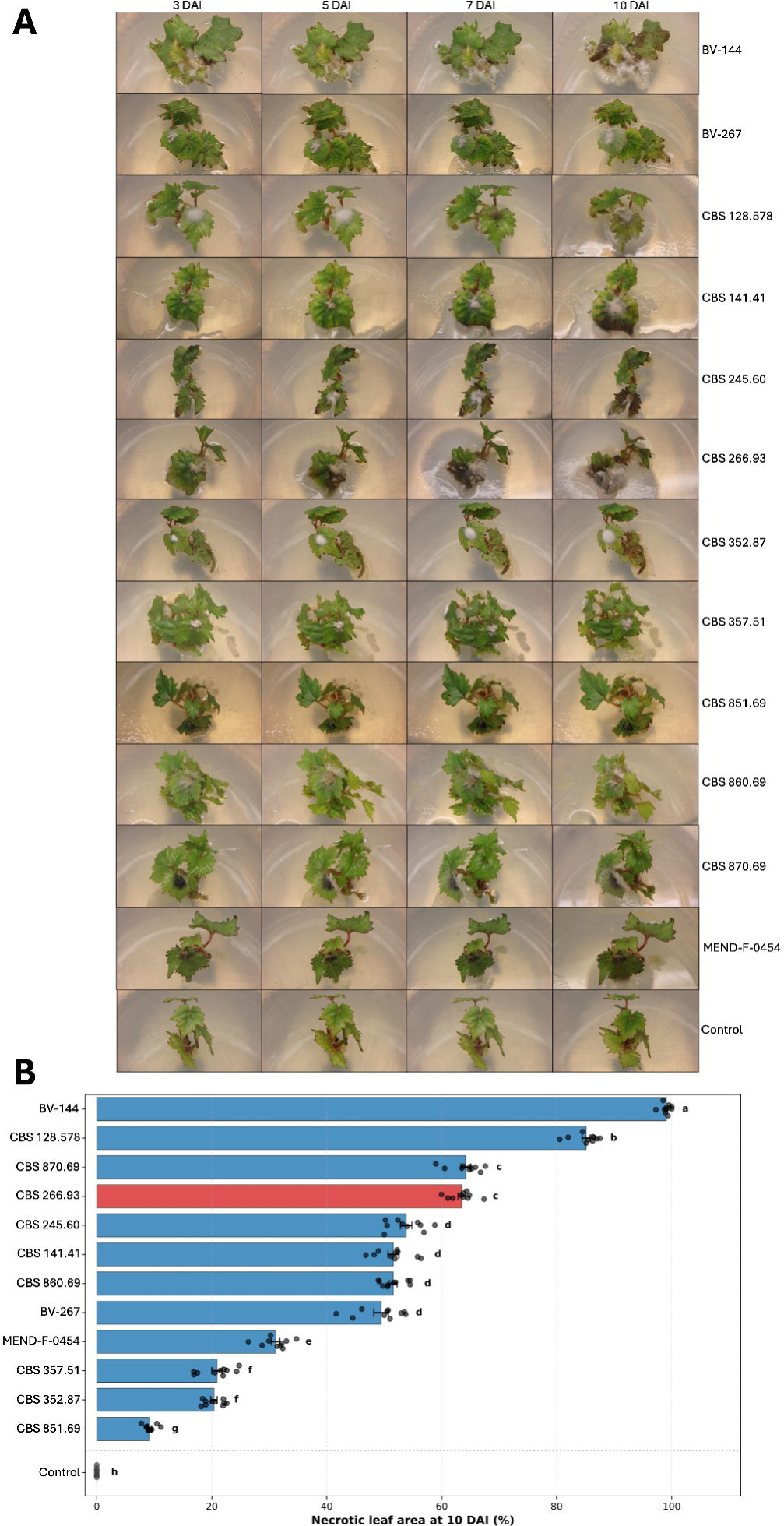

### Plant miRNA Response Analysis

Small RNA profiling revealed marked isolate-associated variation in grapevine miRNA family expression patterns following inoculation with *C. luteo-olivacea* isolates (Figure 8). After abundance filtering and annotation, 70 of 88 retained sequences were assigned with high confidence to known miRNA families, and these were collapsed to family level for downstream analysis. Differential expression analysis relative to the reference isolate BV-144 identified several miRNA families with contrasting patterns across isolates, including miR398, miR827, miR156, miR403, and miR482 (Figure 8A).

**Fig 8.**
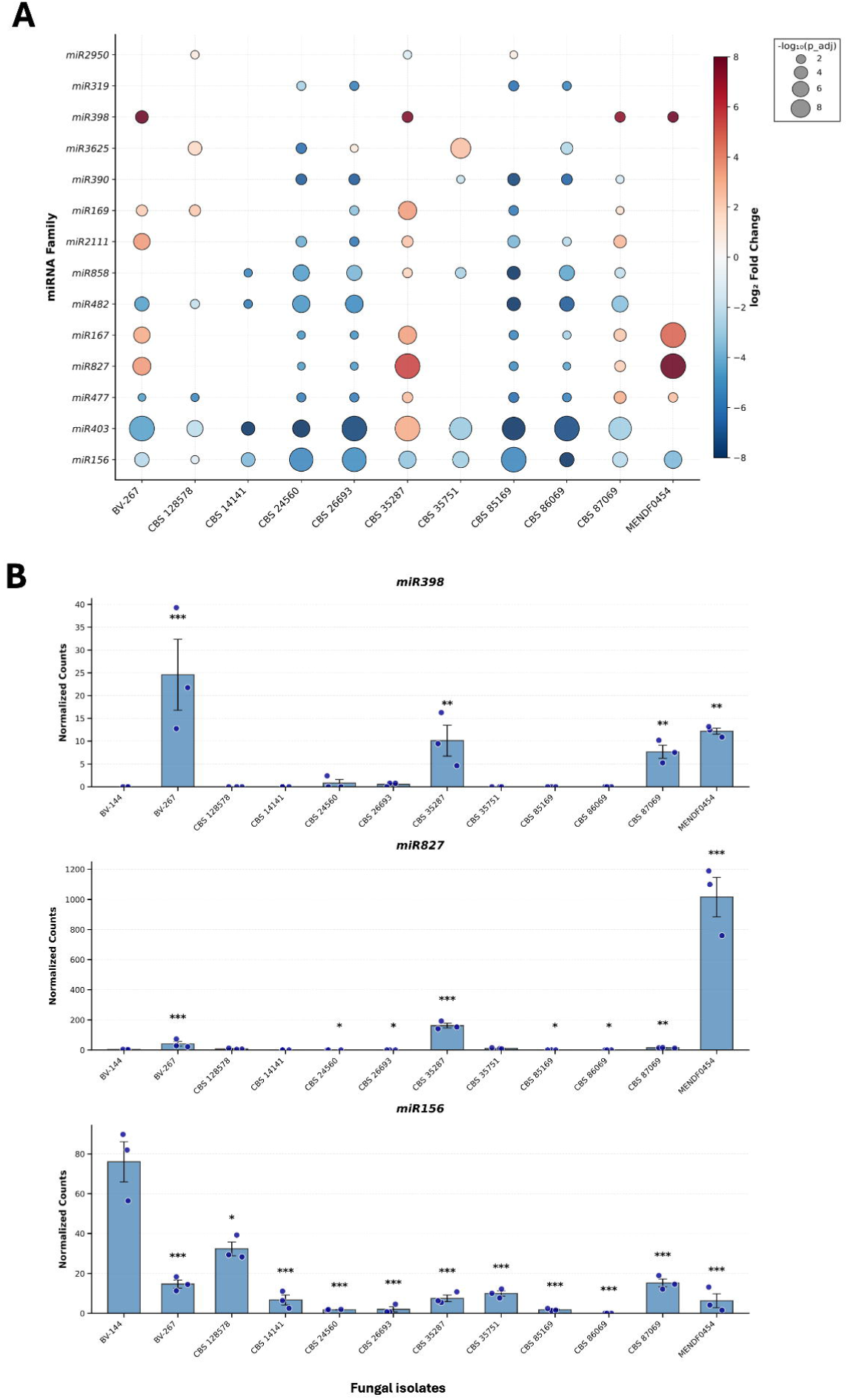

Among the families highlighted in Figure 8B, miR398 showed strongly isolate-dependent accumulation, with highest normalized expression in BV-267 and significantly elevated levels also detected in CBS 352.87, CBS 870.69, and MEND-F-0454 relative to BV-144. In contrast, miR827 displayed a highly asymmetric pattern, with particularly strong accumulation in MEND-F-0454 and, to a lesser extent, in CBS 352.87 and BV-267. miR156 showed the opposite trend, with its highest expression in BV-144 and lower abundance in most of the remaining isolates, indicating a broad shift in this family across the treatments.

At the global level, the bubble plot revealed that different fungal isolates were associated with distinct host miRNA family profiles rather than a single common response pattern (Figure 8A). MEND-F-0454 showed particularly strong positive shifts for miR827 and miR167, whereas several isolates, including CBS 245.60, CBS 266.93, CBS 851.69, and CBS 860.69, were associated with reduced abundance of families such as miR403, miR482, and miR156 relative to BV-144. These results support the existence of substantial intra-specific variation in host-associated miRNA responses among *C. luteo-olivacea* isolates.

Because the main statistical comparisons were performed among inoculated treatments, these miRNA patterns should be interpreted as isolate-associated differences in host small RNA profiles rather than direct infection-versus-control responses. Nevertheless, the marked variation observed for specific families, particularly miR398, miR827, and miR156, suggests that different *C. luteo-olivacea* isolates are associated with distinct regulatory states in grapevine leaves.

## DISCUSSION

This comparative genomic analysis provides a broad framework for understanding genomic diversity, functional conservation, and pathogenicity variation in *C. luteo-olivacea*. Across most of the collection, the species showed a largely cohesive genomic background despite its broad ecological representation, supporting the view that *C. luteo-olivacea* is an adaptable fungus capable of persisting across both plant-associated and environmental contexts. At the same time, the dataset also revealed meaningful lineage-level and phenotypic variation, indicating that this overall cohesion does not imply strict uniformity across strains.

One of the clearest outcomes of this study is the distinct position of strain CBS 266.93. In the ANI analysis, CBS 266.93 consistently showed markedly lower similarity to all other genomes, whereas the remaining 11 strains formed a high-similarity group. The phylogenomic analysis refined this result by showing that CBS 266.93 does not cluster within the principal 11-strain *C. luteo-olivacea* clade, but instead forms a strongly supported sister relationship with *C. malorum*. This pattern suggests that CBS 266.93 is not simply an unusually divergent member of the main lineage, but instead represents a phylogenomically distinct lineage whose taxonomic status should be reassessed. At present, the most conservative interpretation is that CBS 266.93 should be treated as a distinct *Cadophora* lineage pending formal taxonomic reassessment based on expanded taxon sampling and integration of genome-scale, multilocus, and morphological evidence. In this sense, the study not only captures intraspecific diversity, but also highlights the need for continued re-evaluation of species boundaries within *Cadophora*.

Excluding CBS 266.93, the remaining strains showed no clear genomic structuring according to host, environmental origin, or geography. This lack of correspondence between genome-wide similarity and isolation source suggests that diversification within the main *C. luteo-olivacea* lineage has not been driven primarily by simple ecological partitioning. Instead, the pattern is more consistent with a broadly generalist lineage able to persist across multiple substrates while retaining a shared functional backbone. The KOG analysis supports this interpretation, as broad functional categories remained highly similar across strains despite measurable sequence-level divergence. Such decoupling between ecological origin and large-scale functional partitioning is compatible with a flexible lifestyle in which core metabolic and cellular capacities are maintained across multiple habitats (Hill et al., 2022; Deng et al., 2025). More broadly, comparative genome analyses of plant-associated ascomycetes have shown that different pathogenic or host-associated lifestyles can be supported by overlapping virulence-related repertoires rather than by simple ecological or phylogenetic partitioning alone (Wang et al., 2022).

An additional finding relevant to strain identity verification was the ANI = 100% between CBS 128.578 and CBS 141.41. Because the available provenance metadata could not be resolved unambiguously, the relationship between these accessions remains uncertain. These accessions may represent duplicate records, unresolved metadata inconsistencies, or genuinely near-identical isolates. We therefore retained both strains for transparency but interpreted their contribution to broader comparative patterns with caution. Definitive resolution would require raw-read SNP analysis. More broadly, this case underscores the importance of combining genome-scale comparisons with careful metadata curation when using historical culture collection material.

A central feature of this conserved functional framework is the extensive CAZyme repertoire identified across all genomes. The abundance of glycoside hydrolases and auxiliary activity enzymes indicates a broad capacity to degrade plant-derived polysaccharides and modify lignocellulosic substrates, a profile consistent with the colonization of woody tissues (Lombard et al., 2014; Zhao et al., 2013). Moreover, the similarity between *C. luteo-olivacea* CAZyme repertoires and those of established grapevine trunk pathogens suggests that this species shares a functional toolkit compatible with wood colonization and persistence in perennial hosts. This interpretation is also consistent with earlier comparative genomic work on grapevine trunk pathogens showing expansions of gene families associated with plant cell wall degradation, secondary metabolism, and nutrient uptake as recurrent features of these fungi (Morales-Cruz et al., 2015). The widespread presence of salicylate hydrolase genes further supports the idea that all strains retain functions potentially relevant to plant interaction, although the actual contribution of these genes to host colonization remains to be tested experimentally. This point is especially relevant because salicylic acid is a central component of plant defence signalling (Dempsey et al., 2011). Taken together, these results indicate that the capacity to exploit plant structural polymers is not restricted to particular host-associated strains, but instead appears to be a broadly conserved feature of the species. This interpretation is further supported by recent comparative and pangenomic analyses of other grapevine trunk pathogens showing that secreted CAZyme repertoires often remain concentrated in the core functional genome even when other virulence-associated features vary more extensively among lineages (Garcia et al., 2024).

The secondary metabolite results point in the same direction. All strains carried multiple biosynthetic gene clusters, with terpene, PKS, NRPS-like, and NRPS classes dominating across the dataset. Although the total number of clusters varied moderately, their overall composition remained broadly similar, suggesting that the main biosynthetic capacities of *C. luteo-olivacea* are also conserved. At the same time, the fact that many predicted clusters showed little similarity to known pathways suggests considerable unexplored chemical diversity. Rather than indicating strong niche specialization, the current data are more consistent with a species that maintains a versatile and partly cryptic biosynthetic repertoire that may contribute to environmental persistence, fungal competition, and plant-associated survival (Keller, 2019). Functional characterization of these clusters will be needed to determine which metabolites are relevant to pathogenicity and which are more closely tied to saprotrophic or environmental life stages. Recent pangenomic work in other trunk disease fungi has similarly shown that biosynthetic and other virulence-associated functions can be unevenly distributed across variable genomic compartments, even when broad plant-associated fitness is retained at the species level (Garcia et al., 2024).

In contrast to the relative stability of the broader functional profiles, transposable element content varied substantially among strains. Such variation suggests that genome plasticity may differ among lineages even when overall genome size, gene content, and plant-colonizing ability remain similar. Differences in TE load may reflect a combination of historical expansion, silencing efficiency, demographic effects, or lineage-specific exposure to selective pressures favouring either plasticity or genome stability (Muszewska et al., 2019; Fouché et al., 2018). More generally, repeat dynamics are increasingly recognized as a major component of fungal genome plasticity (Sauters and Rokas, 2025). In plant-pathogenic fungi, TEs can contribute to genome restructuring, gene expression variation, and trait evolution (Torres et al., 2021; Abraham et al., 2024), and in some cases large TE insertions have been linked to adaptive traits such as metal resistance (Urquhart et al., 2022). However, TE variation did not map clearly onto aggressiveness, CAZyme content, or the major lineage split identified by phylogenomics, indicating that repeat expansion alone is unlikely to explain the phenotypic differences observed here. This is in line with recent pangenomic analyses of grapevine trunk pathogens in which repeat-rich and variable genomic compartments appear to contribute disproportionately to structural variation, while not necessarily translating into simple genome-wide predictors of aggressiveness (Garcia et al., 2024).

The pathogenicity assays reinforce the view that plant-colonizing ability is broadly retained across *C. luteo-olivacea*. All strains caused lesions on grapevine leaves *in vitro*, but lesion severity differed significantly among isolates, indicating substantial quantitative variation in aggressiveness. These results suggest that the capacity to colonize grapevine tissue is not restricted to a subset of host-associated isolates, although the magnitude of damage varies considerably among strains. Because the assays were performed on leaves of a single cultivar under controlled conditions, these findings should be interpreted as comparative evidence of plant-colonizing ability rather than as a direct proxy for field virulence. Even so, the observation that isolates from non-grapevine or non-plant-associated origins also retained the ability to induce lesions supports the idea that plant-associated fitness is broadly maintained across the species. The observation that some strains produced dark melanized mycelium whereas others did not suggest additional phenotypic variation in traits potentially linked to stress tolerance or survival (Jacobson, 2000). In this context, it is notable that TE insertions have been shown to influence melanin production and associated regulatory variation in other fungal pathogens (Krishnan et al., 2018). However, melanisation was not tightly coupled here to either genome-wide similarity or the highest lesion severity, indicating that pigmentation should not be interpreted as a simple proxy for virulence.

The host small RNA analysis adds a complementary phenotypic dimension to these strain-level differences. Rather than identifying a single uniform host response, the data revealed isolate-associated variation in grapevine miRNA family profiles, particularly involving miR398, miR827, and miR156. Because the final statistical comparisons were performed among inoculated treatments rather than against a well-powered mock control, and because mock libraries showed substantially lower read depth, these patterns should be interpreted cautiously as relative host response states associated with different fungal isolates rather than as canonical infection-induced signatures. Even so, the repeated involvement of families linked in plants to stress signalling and developmental regulation suggests that genetically and phenotypically distinct *C. luteo-olivacea* isolates are associated with different host regulatory states (Sunkar et al., 2012; Li et al., 2022). In this context, the miRNA data are best viewed as hypothesis-generating and as a useful first step towards more targeted functional work on host responses to *Cadophora* colonization.

Overall, the picture that emerges is not one of sharp functional specialization by host or environment, but of a species with a conserved genomic and functional backbone, within which individual lineages differ in phylogenomic placement, repeat content, aggressiveness, pigmentation, and host-associated response profiles. This combination of conservation and variation is likely to be central to the ecological success of *C. luteo-olivacea*. More broadly, recent phylogenomic and pangenomic studies of other grapevine trunk pathogens have shown that important components of virulence-related diversity may only become fully apparent when broader sampling and explicit core-versus-accessory genome analyses are applied (Garcia et al., 2021, 2024). The present study extends that type of comparative framework to *Cadophora* and shows that, in this genus, broad functional conservation can coexist with clear lineage-level divergence and quantitative phenotypic variation. Future work should prioritise the taxonomic reassessment of CBS 266.93, functional characterization of candidate secondary metabolite pathways, and experimental validation of the host regulatory responses highlighted by the miRNA analysis. Such studies will be essential for determining how conserved genomic repertoires translate into the quantitatively different outcomes observed across strains.

## CONCLUSIONS

This study provides the first broad comparative genomic framework for *C. luteo-olivacea* across multiple hosts and environments. Most analysed strains shared a conserved genomic backbone, including similar broad functional profiles, extensive CAZyme repertoires, abundant biosynthetic gene clusters, and widely maintained functions potentially relevant to plant colonization. At the same time, meaningful variation was detected in transposable element content, quantitative aggressiveness, mycelial phenotype, and host-associated miRNA family profiles.

A major finding was the consistent divergence of CBS 266.93, which showed reduced ANI values relative to all other strains and occupied a distinct position in the supplementary phylogenomic analysis, forming a sister relationship with *C. malorum*. This result indicates that the taxonomic placement of CBS 266.93 should be reassessed and highlights the value of genome-scale approaches for refining species boundaries within *Cadophora*.

Despite this divergent lineage, the remaining isolates showed broad conservation of genomic features compatible with plant tissue colonization, supporting the view that *C. luteo-olivacea* retains a shared functional framework across ecologically diverse strains. Quantitative pathogenicity assays further showed that all isolates colonized grapevine leaves in vitro, although lesion severity differed significantly among strains, indicating that plant-colonizing ability is broadly retained but varies in magnitude among isolates.

Finally, host small RNA profiling revealed isolate-associated shifts in grapevine miRNA family expression, especially involving miR398, miR827, and miR156. Although these patterns should be interpreted cautiously as relative differences among inoculated treatments rather than canonical infection-versus-control signatures, they suggest that strain-level diversity in *C. luteo-olivacea* is reflected not only in genome variation but also in distinct host-associated response profiles.

Overall, these findings position *C. luteo-olivacea* as a fungus with a largely conserved genomic and functional background, within which lineage divergence and strain-level phenotypic variation contribute to ecological flexibility and plant-associated behaviour. The results provide a useful basis for future work on taxonomy, secondary metabolism, pathogenicity, and host interaction in *Cadophora*.

## AUTHOR CONTRIBUTIONS

LA performed bioinformatics data processing and analysis. SC analysed genomic data and provided critical manuscript revisions. CL, RB, AE, JP, and EH contributed to experimental design, sample collection, and laboratory analyses. DG conceived and supervised the study, secured funding, and contributed to manuscript writing and editing. All authors reviewed and approved the final manuscript.

## AVAILABILITY OF DATA AND MATERIALS

Raw sequencing reads have been deposited in the NCBI Sequence Read Archive under BioProject PRJNA1430357, with individual SRA accession numbers SRR37421662-SRR37421673. Genome assemblies have been deposited in GenBank under the corresponding assembly accession numbers for each strain. Individual accession numbers for raw reads and genome assemblies are provided in Table 1.

## ACKNOWLEDGMENTS

The authors thank the CBS collection for providing reference strains and acknowledge the technical assistance provided by laboratory staff at all participating institutions. The authors are also grateful to Josep Armengol (Instituto Agroforestal Mediterráneo, Universitat Politècnica de València, Spain) for providing the Spanish isolate Clo-46 from his collection.

## REFERENCES

Abraham, L.N., Oggenfuss, U., Croll, D., 2024. Population-level transposable element expression dynamics influence trait evolution in a fungal crop pathogen. mBio 15, e02840–23.

Aislabie, J., Foght, J., Saul, D., 2001. Aromatic hydrocarbon-degrading bacteria from soil near Scott Base, Antarctica. Polar Biol. 23, 183–188.

Allington, W.B., Chamberlain, D.W., 1948. Brown stem rot of soybean. Phytopathology 38, 793–802.

Andrews, S., 2010. FastQC: a quality control tool for high throughput sequence data.

Arenz, B.E., Held, B.W., Jurgens, J.A., Farrell, R.L., Blanchette, R.A., 2006. Fungal diversity in soils and historic wood from the Ross Sea Region of Antarctica. Soil Biol. Biochem. 38, 3057–3064.

Bankevich, A., Nurk, S., Antipov, D., Gurevich, A.A., Dvorkin, M., Kulikov, A.S., et al., 2012. SPAdes: a new genome assembly algorithm and its applications to single-cell sequencing. J. Comput. Biol. 19, 455–477.

Baránek, M., Křižan, B., Ondrušíková, E., Pidra, M., 2010. DNA-methylation changes in grapevine somaclones following in vitro culture and thermotherapy. Plant Cell Tissue Organ Cult. 101, 11–22.

Bengtsson-Palme, J., Ryberg, M., Hartmann, M., Branco, S., Wang, Z., Godhe, A., et al., 2013. Improved software detection and extraction of ITS1 and ITS2 from ribosomal ITS sequences of fungi and other eukaryotes for analysis of environmental sequencing data. Methods Ecol. Evol. 4, 914–919.

Blanchette, R.A., Held, B.W., Jurgens, J.A., McNew, D.L., Harrington, T.C., Duncan, S.M., Farrell, R.L., 2004. Wood-destroying soft rot fungi in the historic expedition huts of Antarctica. Appl. Environ. Microbiol. 70, 1328–1335.

Blin, K., Shaw, S., Steinke, K., Villebro, R., Ziemert, N., Lee, S.Y., et al., 2019. antiSMASH 5.0: updates to the secondary metabolite genome mining pipeline. Nucleic Acids Res. 47, W81–W87.

Borodovsky, M., Lomsadze, A., 2011. Eukaryotic gene prediction using GeneMark.hmm-E and GeneMark-ES. Curr. Protoc. Bioinformatics Chapter 4, Unit 4.6.10.

Buchfink, B., Xie, C., Huson, D.H., 2015. Fast and sensitive protein alignment using DIAMOND. Nat. Methods 12, 59–60.

Chen, Q., Bakhshi, M., Balci, Y., Broders, K.D., Cheewangkoon, R., Chen, S.F., Fan, X.L., Gramaje, D., Halleen, F., Jung, M.H., Jiang, N., Jung, T., Májek, T., Marincowitz, S., Milenković, I., Mostert, L., Nakashima, C., Nurul Faziha, I., Pan, M., Raza, M., Scanu, B., Spies, C.F.J., Suhaizan, L., Suzuki, H., Tian, C.M., Tomšovský, M., Úrbez-Torres, J.R., Wang, W., Wingfield, B.D., Wingfield, M.J., Yang, Q., Yang, X., Zare, R., Zhao, P., Groenewald, J.Z., Cai, L., Crous, P.W., 2022. Genera of phytopathogenic fungi: GOPHY 4. Stud. Mycol. 101, 417–564.

Chen, S., Zhou, Y., Chen, Y., Gu, J., 2018. fastp: an ultra-fast all-in-one FASTQ preprocessor. Bioinformatics 34, i884–i890.

Darriba, D., Taboada, G.L., Doallo, R., Posada, D., 2012. jModelTest 2: more models, new heuristics and parallel computing. Nat. Methods 9, 772.

Dempsey, D.A., Vlot, A.C., Wildermuth, M.C., Klessig, D.F., 2011. Salicylic acid biosynthesis and metabolism. Arabidopsis Book 9, e0156.

Deng, Z.L., Dissanayake, A.J., Zhu, J.T., et al., 2025. Genomic evolution and diversity in Botryosphaeriales: insights from pan-genomic and population genetic analyses of representative species. Fungal Divers. 134, 1–18.

Fouché, S., Plissonneau, C., Croll, D., 2018. The birth and death of effectors in rapidly evolving filamentous pathogen genomes. Curr. Opin. Microbiol. 46, 34–42.

Frith, M.C., 2011. A new repeat-masking method enables specific detection of homologous sequences. Nucleic Acids Res. 39, e23.

Garcia, J.F., Morales-Cruz, A., Cochetel, N., Minio, A., Figueroa-Balderas, R., Rolshausen, P.E., Baumgartner, K., Cantu, D., 2024. Comparative pangenomic insights into the distinct evolution of virulence factors among grapevine trunk pathogens. Mol. Plant Microbe Interact. 37, 127–142.

Garcia, J.F., Lawrence, D.P., Morales-Cruz, A., Travadon, R., Minio, A., Hernandez-Martinez, R., Rolshausen, P.E., Baumgartner, K., Cantu, D., 2021. Phylogenomics of plant-associated Botryosphaeriaceae species. Front. Microbiol. 12, 652802.

Gramaje, D., Eichmeier, A., 2026. Beyond Koch’s postulates: the pathobiome paradigm in grapevine esca disease. FEMS Microbiol. Ecol. 102, fiag028.

Gramaje, D., León, M., Santana, M., Crous, P.W., Armengol, J., 2014. Multilocus ISSR markers reveal two major genetic groups in Spanish and South African populations of the grapevine fungal pathogen Cadophora luteo-olivacea. PLoS One 9, e110417.

Gramaje, D., Mostert, L., Armengol, J., 2011. Characterization of Cadophora luteo-olivacea and C. melinii isolates obtained from grapevines and environmental samples from grapevine nurseries in Spain. Phytopathol. Mediterr. 50, S112–S126.

Gramaje, D., Úrbez-Torres, J.R., Sosnowski, M.R., 2018. Managing grapevine trunk diseases with respect to etiology and epidemiology: current strategies and future prospects. Plant Dis. 102, 12–39.

Gramaje, D., Mostert, L., Trouillas, F.P., Úrbez-Torres, J.R., Alves, A., 2025. The most informative loci to identify trunk disease pathogens associated with grapevine and perennial fruit and nut crops. Phytopathol. Mediterr. 64, 631–636.

Gurevich, A., Saveliev, V., Vyahhi, N., Tesler, G., 2013. QUAST: quality assessment tool for genome assemblies. Bioinformatics 29, 1072–1075.

Haas, B.J., Salzberg, S.L., Zhu, W., Pertea, M., Allen, J.E., Orvis, J., et al., 2008. Automated eukaryotic gene structure annotation using EVidenceModeler and the Program to Assemble Spliced Alignments. Genome Biol. 9, R7.

Halleen, F., Mostert, L., Crous, P.W., 2007. Pathogenicity testing of lesser-known vascular fungi of grapevines. Australas. Plant Pathol. 36, 277–285.

Held, B.W., Jurgens, J.A., Arenz, B.E., Duncan, S.M., Farrell, R.L., Blanchette, R.A., 2005. Environmental factors influencing microbial growth inside the historic expedition huts of Ross Island, Antarctica. Int. Biodeterior. Biodegrad. 55, 45–53.

Hill, R., Buggs, R.J.A., Toan Vu, D., Gaya, E., 2022. Lifestyle transitions in fusarioid fungi are frequent and lack clear genomic signatures. Mol. Biol. Evol. 39, msac085.

Huerta-Cepas, J., Szklarczyk, D., Heller, D., Hernández-Plaza, A., Forslund, S.K., Cook, H., et al., 2019. eggNOG 5.0: a hierarchical, functionally and phylogenetically annotated orthology resource based on 5090 organisms and 2502 viruses. Nucleic Acids Res. 47, D309–D314.

Hujslová, M., Kubátová, A., Chudíčková, M., Kolařík, M., 2010. Diversity of fungal communities in saline and acidic soils in the Soos National Natural Reserve, Czech Republic. Mycol. Prog. 9, 1–15.

Jacobson, E.S., 2000. Pathogenic roles for fungal melanins. Clin. Microbiol. Rev. 13, 708–717.

Jain, C., Rodriguez-R, L.M., Phillippy, A.M., Konstantinidis, K.T., Aluru, S., 2018. High throughput ANI analysis of 90K prokaryotic genomes reveals clear species boundaries. Nat. Commun. 9, 5114.

Jones, P., Binns, D., Chang, H.Y., Fraser, M., Li, W., McAnulla, C., et al., 2014. InterProScan 5: genome-scale protein function classification. Bioinformatics 30, 1236–1240.

Keller, N.P., 2019. Fungal secondary metabolism: regulation, function and drug discovery. Nat. Rev. Microbiol. 17, 167–180.

Kerry, E., 1990. Microorganisms colonizing plants and soil subjected to different degrees of human activity, including petroleum contamination, in the Vestfold Hills and Mac. Robertson Land, Antarctica. Polar Biol. 10, 423–430.

Korf, I., 2004. Gene finding in novel genomes. BMC Bioinformatics 5, 59.

Krishnan, P., Meile, L., Plissonneau, C., Ma, X., Hartmann, F.E., Croll, D., McDonald, B.A., Sánchez-Vallet, A., 2018. Transposable element insertions shape gene regulation and melanin production in a fungal pathogen of wheat. BMC Biol. 16, 78.

Laetsch, D.R., Blaxter, M.L., 2017. BlobTools: interrogation of genome assemblies. F1000Research 6, 1287.

Lagerberg, T., Lundberg, G., Melin, E., 1927. Biological and practical researches into blueing in pine and spruce. Sven. Skogsvårdsfören. Tidskr. 25, 145–272.

Langmead, B., Salzberg, S.L., 2012. Fast gapped-read alignment with Bowtie 2. Nat. Methods 9, 357–359.

Li, H., 2018. Minimap2: pairwise alignment for nucleotide sequences. Bioinformatics 34, 3094–3100.

Li, J., Song, Q., Zuo, Z.F., Liu, L., 2022. MicroRNA398: a master regulator of plant development and stress responses. Int. J. Mol. Sci. 23, 10803.

Lombard, V., Golaconda Ramulu, H., Drula, E., Coutinho, P.M., Henrissat, B., 2014. The carbohydrate-active enzymes database (CAZy) in 2013. Nucleic Acids Res. 42, D490–D495.

Lowe, T.M., Eddy, S.R., 1997. tRNAscan-SE: a program for improved detection of transfer RNA genes in genomic sequence. Nucleic Acids Res. 25, 955–964.

Majoros, W.H., Pertea, M., Salzberg, S.L., 2004. TigrScan and GlimmerHMM: two open source ab initio eukaryotic gene-finders. Bioinformatics 20, 2878–2879.

Marchese, P., Garzoli, L., Young, R., Allcock, L., Barry, F., Tuohy, M., Murphy, M., 2021. Fungi populate deep-sea coral gardens as well as marine sediments in the Irish Atlantic Ocean. Environ. Microbiol. 23, 4168–4184.

Morales-Cruz, A., Amrine, K.C.H., Blanco-Ulate, B., Lawrence, D.P., Travadon, R., Rolshausen, P.E., Baumgartner, K., Cantu, D., 2015. Distinctive expansion of gene families associated with plant cell wall degradation, secondary metabolism, and nutrient uptake in the genomes of grapevine trunk pathogens. BMC Genomics 16, 469.

Morrell, J.J., Zabel, R.A., 1985. Wood strength and weight loss caused by soft-rot fungi isolated from treated southern pine utility poles. Wood Fiber Sci. 17, 132–143.

Muszewska, A., Steczkiewicz, K., Stepniewska-Dziubinska, M.M., Ginalski, K., 2019. Transposable elements contribute to fungal genes and impact fungal lifestyle. Sci. Rep. 9, 4307.

Nilsson, R.H., Larsson, K.H., Taylor, A.F.S., Bengtsson-Palme, J., Jeppesen, T.S., Schigel, D., et al., 2019. The UNITE database for molecular identification of fungi: handling dark taxa and parallel taxonomic classifications. Nucleic Acids Res. 47, D259–D264.

Nilsson, T., 1973. Studies on degradation and cellulolytic activity of microfungi. Stud. For. Suec. 104, 1–40.

Okonechnikov, K., Conesa, A., García-Alcalde, F., 2016. Qualimap 2: advanced multi-sample quality control for high-throughput sequencing data. Bioinformatics 32, 292–294.

Ou, S., Su, W., Liao, Y., Chougule, K., Agda, J.R., Hellinga, A.J., et al., 2019. Benchmarking transposable element annotation methods for creation of a streamlined, comprehensive pipeline. Genome Biol. 20, 275.

Palmer, J.M., Stajich, J.E., 2020. Funannotate v1.8.1: eukaryotic genome annotation. Zenodo.

Prodi, A., Sandalo, S., Tonti, S., Nipoti, P., Pisi, A., 2008. Phialophora-like fungi associated with kiwifruit elephantiasis. J. Plant Pathol. 90, 487–494.

Rouxel, T., Balesdent, M.H., 2017. Life, death and rebirth of avirulence effectors in a fungal pathogen. New Phytol. 214, 526–562.

Sánchez-Vallet, A., Fouché, S., Fudal, I., Hartmann, F.E., Soyer, J.L., Tellier, A., Croll, D., 2018. The genome biology of effector gene evolution in filamentous plant pathogens. Annu. Rev. Phytopathol. 56, 21–40.

Sauters, T.J.C., Rokas, A., 2025. Patterns and mechanisms of fungal genome plasticity. Curr. Biol. 35, R527–R544.

Simão, F.A., Waterhouse, R.M., Ioannidis, P., Kriventseva, E.V., Zdobnov, E.M., 2015. BUSCO: assessing genome assembly and annotation completeness with single-copy orthologs. Bioinformatics 31, 3210–3212.

Slater, G.S.C., Birney, E., 2005. Automated generation of heuristics for biological sequence comparison. BMC Bioinformatics 6, 31.

Stanke, M., Keller, O., Gunduz, I., Hayes, A., Waack, S., Morgenstern, B., 2006. AUGUSTUS: ab initio prediction of alternative transcripts. Nucleic Acids Res. 34, W435–W439.

Sunkar, R., Li, Y.F., Jagadeeswaran, G., 2012. Functions of microRNAs in plant stress responses. Trends Plant Sci. 17, 196–203.

Tamura, K., Stecher, G., Peterson, D., Filipski, A., Kumar, S., 2013. MEGA6: molecular evolutionary genetics analysis version 6.0. Mol. Biol. Evol. 30, 2725–2729.

The UniProt Consortium, 2019. UniProt: a worldwide hub of protein knowledge. Nucleic Acids Res. 47, D506–D515.

Torres, D.E., Thomma, B.P.H.J., Seidl, M.F., 2021. Transposable elements contribute to genome dynamics and gene expression variation in the fungal plant pathogen Verticillium dahliae. Genome Biol. Evol. 13, evab135.

Travadon, R., Lawrence, D.P., Moyer, M.M., Fujiyoshi, P.T., Baumgartner, K., 2022. Fungal species associated with grapevine trunk diseases in Washington wine grapes and California table grapes, with novelties in the genera Cadophora, Cytospora, and Sporocadus. Front. Fungal Biol. 3, 1018140.

Travadon, R., Lawrence, D.P., Rooney-Latham, S., Gubler, W.D., Wilcox, W.F., Rolshausen, P.E., Baumgartner, K., 2015. Cadophora species associated with wood-decay of grapevine in North America. Fungal Biol. 119, 53–66.

Úrbez-Torres, J.R., Haag, P., Bowen, P., O’Gorman, D.T., 2014. Grapevine trunk diseases in British Columbia: incidence and characterization of the fungal pathogens associated with esca and Petri diseases of grapevine. Plant Dis. 98, 469–482.

Urquhart, A.S., Chong, N.F., Yang, Y., Idnurm, A., 2022. A large transposable element mediates metal resistance in the fungus Paecilomyces variotii. Curr. Biol. 32, 937–950.e5.

Wang, Y., Wu, J., Yan, J., Guo, M., Xu, L., Hou, L., Zou, Q., 2022. Comparative genome analysis of plant ascomycete fungal pathogens with different lifestyles reveals distinctive virulence strategies. BMC Genomics 23, 34.

Zhang, H., Yohe, T., Huang, L., Entwistle, S., Wu, P., Yang, Z., et al., 2018. dbCAN2: a meta server for automated carbohydrate-active enzyme annotation. Nucleic Acids Res. 46, W95–W101.

Zhao, Z., Liu, H., Wang, C., Xu, J.R., 2013. Comparative analysis of fungal genomes reveals different plant cell wall degrading capacity in fungi. BMC Genomics 14, 274.

